# Short-term perceptual re-weighting in suprasegmental categorization

**DOI:** 10.1101/2021.01.18.427088

**Authors:** Kyle Jasmin, Adam Tierney, Chisom Obasih, Lori Holt

**Author notes:** Correspondence to: Kyle Jasmin, Department of Psychology, Wolfson Building, Royal Holloway, University of London, Egham, Surrey, TW20 0EX. equal contributions (Kyle Jasmin, Adam Tierney).

## Abstract

Segmental speech units such as phonemes are cued by multiple acoustic dimensions (e.g. F0 and duration), but dimensions do not carry equal perceptual weight. The relative perceptual weights of acoustic speech dimensions are not fixed but vary with context. For example, when speech is altered to create an ‘accent’ in which two acoustic dimensions are correlated in a manner opposite that of long-term experience, the dimension that carries less perceptual weight is down-weighted to contribute less in category decisions. It remains unclear, however, whether this short-term reweighting is limited to segmental categorization, or if it extends to categorization of suprasegmental features which span multiple phonemes, syllables, or words, which would suggest that such “dimension-based statistical learning” is a widespread phenomenon in speech perception. Here we investigated the relative contribution of two acoustic dimensions to word emphasis. Participants categorized instances of a two-word phrase pronounced with typical covariation of fundamental frequency (F0) and duration, and in the context of an artificial ‘accent’ in which F0 and duration (established in prior research on English speech as ‘primary’ and ‘secondary’ dimensions, respectively) covaried atypically. When categorizing ‘accented’ speech, listeners rapidly down-weighted the secondary dimension (duration) while continuing to rely on the primary dimension (F0). This result indicates that listeners continually track short-term regularities across speech input and dynamically adjust the weight of acoustic evidence for suprasegmental categories. Thus, dimension-based statistical learning appears to be a widespread phenomenon in speech perception extending to both segmental and suprasegmental categorization.

---

A central challenge in the study of speech communication is understanding how continuous variation across multiple acoustic dimensions is mapped onto discrete categories. Segmental speech categories such as phonemes are not signalled by any single acoustic dimension. Instead, phonemes are conveyed by multiple acoustic dimensions that vary in their diagnosticity or ‘perceptual weight’ in signalling a speech category (Holt & Lotto, 2006; Toscano and McMurray 2010). For example, in clear speech, the phoneme /b/ (as in ‘bat’) is distinguished from /p/ (as in ‘pat’) in part by the time elapsed between the acoustic burst created by release of the articulators and the onset of the periodic signal associated with the vibration of the vocal folds, an interval referred to as the ‘voice onset time’ or VOT (Lisker 1957), which is longer for /p/ than /b/. Whereas VOT is the most reliable cue signalling /b/–/p/ categorization in clear speech for English listeners, at least 16 other acoustic dimensions also contribute, such as the fundamental frequency (F0) at VOT offset and the length of delay in the onset of the first formant (Lisker 1986). Thus, multiple acoustic dimensions contribute to segmental speech categorization, but the diagnosticity of these dimensions in signalling segmental speech categories varies; dimensions carry different *perceptual weight*.

Perceptual weights of acoustic dimensions are context-dependent: when listeners encounter short-term changes in the ways in which dimensions are associated with categories, perceptual weights adjust in response. For example, as we note above, VOT typically carries greater perceptual weight than F0 in signalling /b/ versus /p/ speech categories—when listening conditions are good. However, in the presence of background noise, this pattern reverses, and F0 is more reliably associated with listeners’ categorization responses than VOT (Winn et al., 2013; Holt et al., 2018; Wu & Holt, under review). Perceptual weights of acoustic dimensions also rapidly shift in response to short-term changes in the distribution of acoustic cues experienced in speech input, such as in encountering a talker with an accent. For example, in English, VOT and F0 typically covary such that longer VOT and higher F0 co-occur and signal /p/, whereas shorter VOT and lower F0 frequencies co-occur and signal /b/. When listeners are exposed to an artificial ‘accent’ for which the relationship between F0 and VOT is reversed (e.g. longer VOTs co-occurring with lower F0), they rapidly down-weight reliance on the secondary dimension such that F0 is no longer an effective signal of /b/ versus /p/ category membership (Idemaru & Holt, 2011; 2014; Zhang & Holt, 2018; Schertz et al., 2016; Lehet & Holt, 2017; Wu & Holt, under review). When the short-term input regularities return to English norms, the perceptual weight of F0 quickly returns to baseline levels such that it signals /b/ and /p/ differentially.

This rapid adaptation of how acoustic input dimensions contribute to segmental speech perception is referred to as ‘dimension-based statistical learning’ (Idemaru & Holt, 2011). Importantly, when co-occurrence statistics between two dimensions are reversed (e.g. to create the ‘accent’ described above), the primary dimension (i.e. the one that typically carries greater perceptual weight) continues to signal category membership unambiguously. For instance, since English listeners perceptually weight VOT more than F0 in clear speech, VOT continues to signal /b/-/p/ category membership even in the context of a reversal of the VOT×F0 correlation in short-term speech input that conveys an accent.

Similar to segmental categories, suprasegmental categories are also signalled by multiple, correlated acoustic dimensions. For example, acoustic measurements indicate that word emphasis in English is most strongly correlated with a sudden change in F0, but duration and amplitude are also correlated to lesser degrees (Breen et al., 2010). Evidence suggests that listeners integrate evidence across acoustic dimensions when making decisions about the presence of suprasegmental categories. For example, English listeners are influenced by both pitch and duration patterns when deciding how a sentence should be parsed (Streeter, 1978; Beach, 1991), by pitch, intensity, duration, and vowel quality when deciding whether a syllable is stressed or un-stressed (Chrabaszcz et al., 2014), and by pitch, duration, and intensity when deciding on the location of contrastive focus (i.e., word emphasis; Jasmin et al., 2020).

The dimension-based statistical learning literature establishes that listeners track short-term input regularities and that these regularities dynamically impact the effectiveness of acoustic dimensions in signaling segmental speech categories. But, we do not yet know whether this phenomenon is limited to segmental categorization or is a broader phenomenon. That suprasegmental categories, like segmental categories, are tied to variation across multiple dimensions suggests that disrupting the canonical co-occurrence pattern of acoustic dimensions may cause listeners to down-weight secondary dimensions in signalling suprasegmental category identity. In the case of word emphasis in English, for which F0 is the primary dimension, we expect listeners to down-weight duration (a secondary dimension; Jasmin et al., 2020) upon introduction of an artificial accent that reverses the canonical F0xDuration covariation in English. Prior research indicates that overall weighting of prosodic information as a cue to speaker intent can change in the short term due to context; for example, exposure to a speaker with idiosyncratic/unreliable use of pitch accents leads listeners to down-weight intonation as a cue to speaker intent (Roettger and Franke 2019, Roettger and Rimland 2020). Moreover, when ambiguous pitch contours are consistently paired with a particular interpretation, categorization functions shift according to whether a statement had a positive or a negative interpretation (Kurumada et al. 2018). Here we ask whether the relative weighting of cues to suprasegmental categorization from different acoustic dimensions can also be modulated in the short-term by context.

We presented participants with spoken phrases drawn from a two-dimensional stimulus space in which stimuli varied in the extent to which F0 contours and duration patterns implied emphasis on one of two words (i.e. ‘STUDY music’ vs. ‘study MUSIC’). In the Canonical condition, F0 and duration cues co-occurred in a manner typical of English, whereas in the Accented condition, F0 and duration cues were presented with an ‘accent’ that reversed the canonical co-occurrence patterns of English. In each condition, a minority of trials were test stimuli for which F0 contour was constant and perceptually ambiguous while duration varied to potentially signal emphasis on one word or another. If the perceptual weight of these dimensions is flexibly adjusted according to short-term speech input regularities, then we predict that the test stimuli varying in duration will differentially signal word emphasis (as *STUDY music* vs. *study MUSIC*) in the Canonical condition and be down-weighted to have less influence on suprasegmental categorization in the Accented condition that reverses the canonical dimension relationship.

## Methods

### Participants

Native speakers of American English (N = 43, 37 F; aged 18–22 years) with normal hearing were recruited from Carnegie Mellon University. Participants took part for university credit or payment after giving informed consent. The study was approved by the Carnegie Mellon University Institutional Review Board in line with the Declaration of Helsinki. Based on robust behavioral effects from prior studies of dimension-based statistical learning for segmental speech (e.g., Idemaru & Holt, 2011; Liu & Holt, 2015) which yielded very large effect sizes (e.g. Cohen’s *d =* 1.7 for the analogous interaction effect in a similar study from Liu & Holt, 2015), we estimated that a minimum of 21 participants would be required to detect an effect with predicted power of 99% (conducted using ‘pwr’ package in R, two-tailed alpha at 0.05). However, because dimension-based statistical learning paradigms had not been used previously to examine a suprasegmental contrast, final sample size was determined by collecting the maximum number of participants that could be recruited and tested given time and resource constraints.

### Stimulus Creation

The stimulus space was defined by orthogonal acoustic manipulations across duration and F0 contour over tokens of the spoken English phrase “study music.” The tokens were created by recording the voice of a native English speaker saying the phrases “Dave likes to STUDY music” (early focus) and “Dave likes to study MUSIC” (late focus), with emphasis placed either on STUDY, or MUSIC. The two recordings were then ‘morphed’ together using STRAIGHT software (Kawahara & Irino, 2005; Jasmin et al., 2020ab,c): the F0 was extracted from voiced segments of the two utterances; next, aperiodic aspects of the signal were identified and analyzed; then, the filter characteristics of the signal were calculated. Finally, the two “morphing substrates” (speech from each recording decomposed into F0, aperiodic aspects, and filter characteristics) were manually time aligned by marking corresponding ‘anchor points’ in both recordings. This was done by examining a similarity matrix generated by STRAIGHT (based on the two input sound files) and manually marking corresponding salient changes in the spectrograms.

Following temporal alignment, STRAIGHT’s morphing procedure involves regenerating a signal using a linear interpolation between the manually-marked anchor points in an abstract distance space (Kawahara & Irino, 2005). For F0 this is in the log–frequency domain. In creating these morphed versions, the F0 contour and durational morphing rates were adjusted orthogonally in order to create a 7-by-7 grid of stimuli whose F0 and durational properties cued emphasis on STUDY or on MUSIC to 7 different degrees: 0%, 17%, 33%, 50%, 67%, 83%, and 100%, with 0% indicating that the F0 contour or duration characteristics came from the “STUDY music” recording, 100% meaning F0 and duration were identical to the “study MUSIC” recording, and intermediate values indicating F0 and duration patterns linearly interpolated between the two original recordings. Finally, the stimuli were trimmed to only contain the two words “study” and “music.” Following morphing, the differences in F0 between study and music, measured at the nucleus of the first vowel of each word, at each of the seven F0 levels were –8.5, –5.0, –2.1, +0.6, +3.4 +5.7, and +8.1 semitones, negative values reflecting higher frequency F0 on “music” than “study”. (These steps were not exactly evenly spaced as they reflect the difference in measurements between the two words for F0.) The differences in duration between the first (stressed) syllables of ‘study’ and ‘music’ (measured as the onset and vowel, ‘stuh’ and ‘myoo’) in the final morphed stimuli were approximately 220, 150, 110, 90, 70, 50, and 40, milliseconds. The boundary between syllables was defined by visually inspecting the spectrogram and marking the offset of voicing before the /d/ in ‘study’ and the onset of high frequency fricative information in ‘music’.

### Baseline Stimuli

Figure 1 illustrates how stimuli were sampled from this 7-by-7 stimulus space across blocks. Baseline stimuli consisted of 25 versions of the spoken phrase “study music” with word emphasis manipulated across F0 contour and duration. A 5-by-5 subset from the center of the 7- by-7 stimulus space (grey in Figure 1) sampled the two acoustic dimensions orthogonally to establish listeners’ baseline perceptual weights in labeling the speech as having early versus late word emphasis (*STUDY music* versus *study MUSIC*).

**Figure 1.**
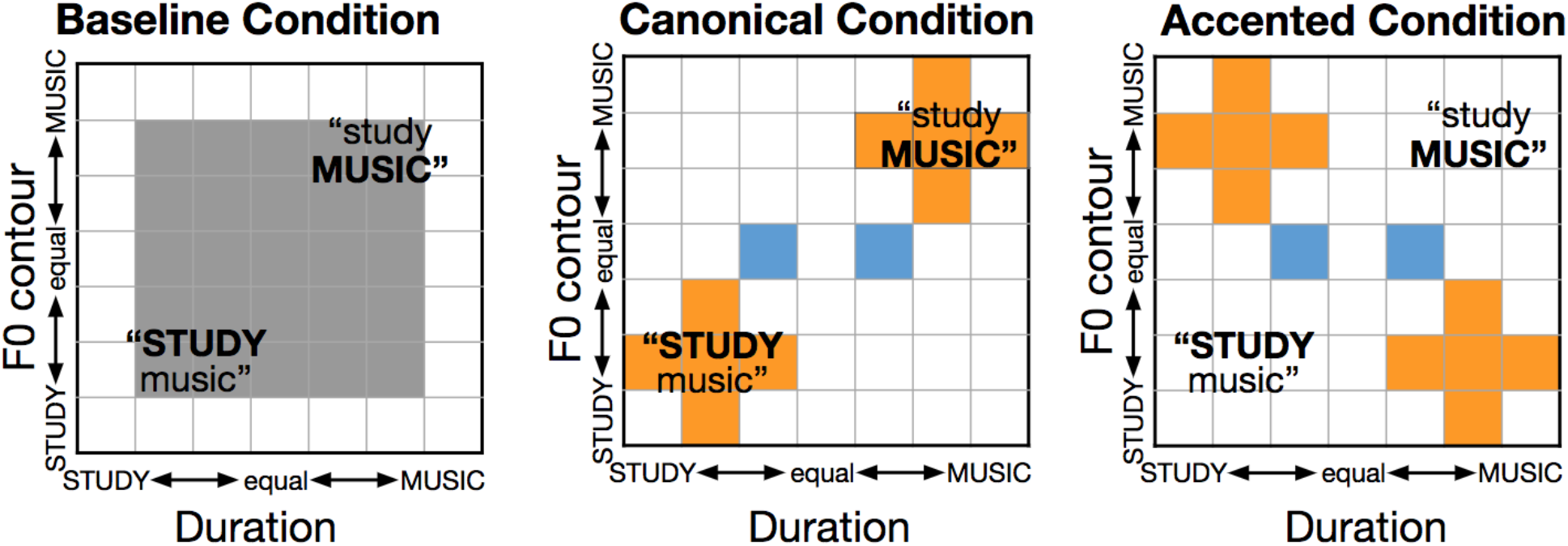
Stimuli. Stimuli sampled a 7-by-7 acoustic space across duration and F0 contour. Baseline categorization measurements made use of the center 25 stimuli in the grid (left panel), sampled orthogonally across dimensions. During the Canonical Block (middle panel), participants categorized canonical-exposure stimuli (orange squares). During accented exposure (right panel), participants categorized stimuli for which F0 and duration cues possessed a correlation opposite that of English (orange squares). Participants also categorized Test stimuli, which had identical F0 contours and distinct durations (blue squares).

### Exposure Stimuli

In subsequent blocks, stimuli were sampled from the 7-by-7 stimulus space to manipulate short-term speech input regularities across a Canonical block that mirrored English acoustic regularities (orange squares, Figure 1) and an Accented block that reversed the typical correlation between F0 contour and duration to create an artificial ‘accent’ (orange squares, Figure 1). Exposure stimuli comprised 80% of trials in these blocks.

### Test Stimuli

Test stimuli (blue squares, Figure 1) made up the remaining 20% of stimuli within the Canonical and Accented blocks. Test stimuli were constant across blocks and served to assess the degree to which listeners make use of duration to signal word emphasis as a function of the short-term regularities conveyed by the Exposure stimuli that vary across blocks. Test stimuli possessed acoustically ambiguous F0 Contour (level 50%) and distinct duration (level 33%, level 67%). Test stimuli were randomly interspersed with Exposure stimuli in the Canonical and Accented blocks.

### Procedure

Participants were seated in front of a computer monitor in a sound-attenuated booth. Each trial began with a looming checkerboard circle in the center of the monitor. When participants had fixated on the checkerboard for one second, a stimulus phrase “study music” (Fig. 1) was presented diotically over headphones (Beyer DT-150) and then the response options appeared on the screen. Participants were instructed to press either the ‘z’ or ‘m’ key on the keyboard, associated with the spatial position of the response labels, to indicate whether they heard “STUDY music” or “study MUSIC.” The key press triggered the next trial.

Participants experienced the Baseline, Canonical and Accented conditions as three blocks, always presented in the same order. All that differed across blocks was the sampling of stimuli. The task remained constant. Trials were presented across blocks without breaks or any other overt demarcation so that block structure was implicit and unknown to participants. The Baseline block consisted of 200 trials (25 stimuli × 8 presentations; grey in Figure 1), the Canonical block consisted of 80 Canonical exposure trials (10 stimuli with 8 presentations; orange, Figure 1, middle panel) and 16 interspersed Test trials (2 stimuli with 8 presentations; blue, Figure 1), and the Accented block consisted of 80 Accented exposure trials (orange, Figure 1, right panel) and 16 interspersed test trials (blue, Figure 1). The entire session was completed in approximately 25 minutes. The experiment was delivered under the control of E-prime experiment software (Psychology Software Tools, Inc.).

### Analyses

F0 contour and duration perceptual weights for the baseline trials were calculated by estimating a logistic regression for each subject, with F0 level (2 to 6) and duration level (2 to 6) predicting the binary response (*STUDY music* vs *study MUSIC*). The coefficients for F0 contour and duration were then combined by normalizing them such that they summed to one (Holt & Lotto 2006; Idemaru et al., 2012; Jasmin et al., 2020a), resulting in a normalized perceptual weight that ranged between 0 and 1, with values closer to 1 indicating greater reliance on F0 contour than duration, values closer to 0 indicating the reverse, and 0.5 indicating equal reliance. The mean normalized perceptual weights were compared across subjects against a value of 0.5 with a one sample t-test.

Performance on the exposure trials in the Canonical and Accented blocks were assessed for accuracy as proportion correct (defined according to the ‘dominant,’ heavily perceptually weighted, dimension from the baseline weights). To analyze effects of Canonical and Accented exposure on categorization of test stimuli, the trial-wise data for all participants was entered into a mixed effects logistic regression using *lme4*’s *glmer* function (Bates et al., 2015) with ‘family=binomial’, and Response (*STUDY music* vs *study MUSIC*) predicted by the Exposure Type (Canonical vs Accented), duration (longer *STUDY* vs longer *MUSIC*), and their interaction, as well as Participant as a random intercept in R. The effect of the interaction term was calculated by comparing this full model to a null model (without the interaction) using R’s *anova* function. Pairwise tests were conducted with *lsmeans* in R (Russell & Length, 2016).

## Results

### Baseline Categorization

Figure 2 illustrates average categorization responses for the Baseline block in which F0 contour and duration varied orthogonally across stimuli. Participants tended to rely more on F0 contour than duration to categorize the spoken phrase according to word emphasis, replicating the results of Jasmin et al. (2020a), and confirming that F0 contour is a stronger cue to word emphasis than duration in English (normalized perceptual cue weight M_F0_ = 0.81±0.03, t(42) = 10.62, *p* < 0.001; higher values indicate greater F0 contour reliance).

**Figure 2.**
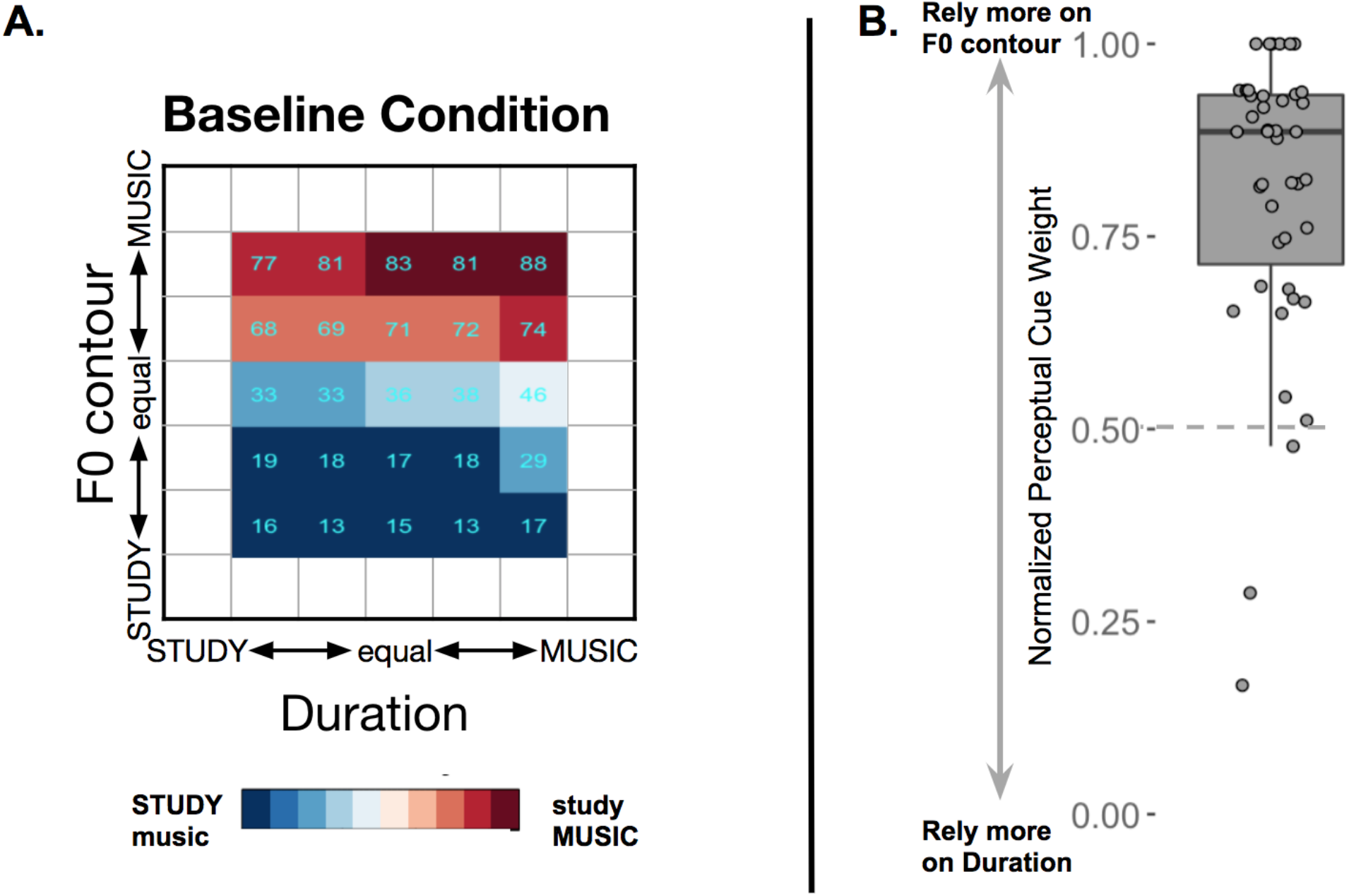
Results from the Baseline Condition. **A)** Mean percent categorization responses for each of the baseline stimuli. Blue indicates that participants tended to perceive emphasis as “*STUDY music*” whereas red indicates that they perceived emphasis as “*study MUSIC*”. **B)** Normalized perceptual weights for each participant. Most participants relied more on the F0 contour dimension than the duration dimension to judge emphasis.

### Categorization of Exposure Stimuli

Responses to the unambiguous exposure stimuli were examined to ensure that participants were using F0 (the primary dimension signalling word emphasis) to make their judgments in the Canonical and Accented blocks (orange squares in Fig.1). The mean percentage of correct responses, defined according to F0 contour, was high during the Canonical block (M = 88.1±12.1) as well as during the Accented block (M=81.9 ± 0.17.2).

### Categorization of Test Stimuli

Test stimuli served as the primary measure of whether short-term speech input regularities impact perception of word emphasis. Recall that test stimuli possessed an acoustically ambiguous F0 contour, thereby neutralizing the acoustic dimension most listeners rely upon to make word emphasis judgements (Figure 2). Thus, categorization of Test stimuli provides a measure of the extent to which listeners rely on duration to judge word emphasis, and whether the perceptual weight of duration is modulated across manipulations in short-term speech regularities experienced across Exposure stimuli in the Canonical and Accented blocks. Figure 3 illustrates these results.

**Figure 3:**
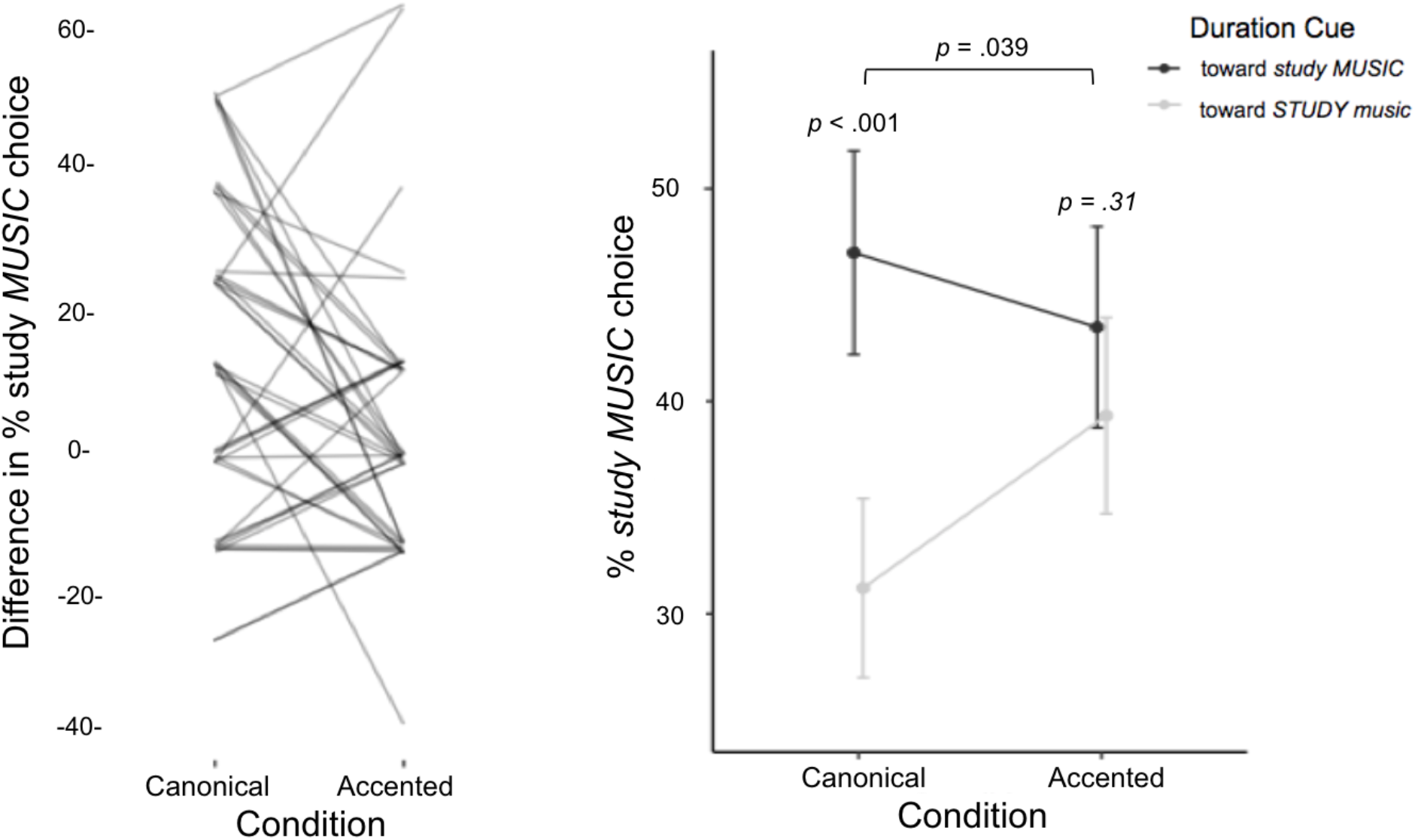
Test Stimulus Categorization in the context of Canonical and Reversed Regularities. Suprasegmental categorization behavior in the context of exposure to Canonical and Accented statistical co-occurrence of F0 contour and duration dimensions. When short-term regularity aligned with long-term English regularities in the Canonical block, duration differentially signalled word emphasis as *STUDY music* versus *study MUSIC*. Nonetheless, categorization of the same stimuli differed when short-term regularities departed from English in the Accented block; participants no longer relied upon the duration dimension in word emphasis judgments. The left panel shows subject-level data: the difference in percent of *study MUSIC* responses across the test trials for the Canonical versus Accented blocks. The right panel shows the mean percentage of responses categorized as *study MUSIC* for each test stimulus individually, and standard errors. Inferential statistics are the results of the mixed model analysis reported in the main text.

As predicted, categorization of the test trials differed as a function of the short-term speech input regularities experienced across the Canonical and Accented blocks (comparison of the full model including interaction of Block and Test Stimulus Duration term with null model omitting the interaction χ^2^(1, 5) = 4.27, p = 0.039). Pairwise post-hoc tests indicated that in the context of Canonical short-term regularities in speech input conveyed by the exposure trials, duration influenced categorization when F0 contour was ambiguous, with longer word duration indicating emphasis (OR = 1.95, Cohen’s *d =* 0.37, Z = 3.90, *p* < .001). However, upon introduction of the artificial ‘accent’ that reversed the relationship between F0 contour and duration relative to canonical English patterns, listeners’ reliance on duration to signal word emphasis rapidly shifted. In the context of exposure to the Accented short-term regularity conveyed by exposure trials the perceptual weight of durational information dramatically decreased, to the point that there was no significant difference in participants’ word emphasis judgements as a function of duration (OR = 1.19, Cohen’s *d =* 0.10, Z = 1.02, *p* = 0.31).

## Discussion

While it has been well-established that the relative weighting of different acoustic dimensions for segmental speech categorization can shift rapidly according to the listening context, it has not previously been established whether similar processes apply to categorization of linguistic features that span multiple segments. In the present work we exposed listeners to an artificial ‘accent’ in which the typical co-occurrence of F0 contour and duration for word emphasis in English was reversed. We found that the perceptual weight of duration sharply decreased in response to this shift in context, and we therefore conclude that perceptual cue weights for prosodic categories are malleable, responding dynamically to statistical properties of the speech input.

In this study we examined a single context manipulation (‘typical’ speech for which F0 and duration cues covaried normally and ‘accented’ speech for which F0 and duration cues were opposite of the typical covariation pattern). We also presented only a single spoken token (“study music”) spoken by a single talker. It remains to be seen, therefore, whether short-term changes in suprasegmental cue weights generalize to other suprasegmental categories, other specific examples of word emphasis, or other speech from other talkers, who may use prosody variably (Peppé et al., 2000). Research on dimension-based statistical learning here has observed that some generalization takes place, but to different extents depending on speaker and linguistic contexts (Idemaru & Holt, 2014; Liu & Holt, 2015; Zhang and Holt, 2018; Lehet and Holt, 2020). For segmental speech perception, it has been shown that learning of an artificial accent does indeed generalize across linguistic contexts (e.g., to lists of words/nonwords (Idemaru and Holt, 2020; Lehet and Holt, 2020) and across voices (Liu & Holt, 2015), but the degree of down-weighting tends to be lesser in contexts not directly experienced by listeners. There is also evidence that speaker information cued vocally or visually can be used to guide speaker-specific dimension-based statistical learning of phonetic categories, supporting even simultaneous tracking of multiple input regularities (Zhang and Holt, 2018). Based on this prior evidence from research on segmental categorization, we predict that suprasegmental down-weighting will generalize across voices but do not have a strong prediction regarding whether down-weighting will generalize to other suprasegmental features (for example, from word emphasis to phrase boundaries).

Though there remain many open questions even for adaptive plasticity across segmental speech categorization, in general patterns of generalization seem to be consistent with an account that category activation drives reweighting (Idemaru & Holt, 2011; Zhang et al., 2021; Wu & Holt, under review; Wu, 2020) and the mechanism for this may involve supervised error-driven learning (Guediche et al., 2014; Wu, 2000) or reinforcement learning (Harmon et al., 2019). To the extent that short-term input regularities across acoustic dimensions are effective in activating a suprasegmental category even as they deviate from long-term expectations of correlations among input dimensions, we would anticipate re-weighting and modest generalization. In fact, as has been the case in studies of segmental categories (Idemaru and Holt, 2014), successes and failures of the generalization of dimension-based statistical learning can inform the nature of underlying category representations. The present results suggest that dimension-based statistical learning could also be used to investigate the nature of the representations underlying suprasegmental categories.

It remains unknown whether other prosodic features signalled by multiple acoustic dimensions are subject to dimension-based statistical learning. Further work could examine, for example, intonational phrase boundaries, which are accompanied acoustically by lengthening of the syllable just before the boundary and increased pause duration (Choi et al. 2005, Cumming 2010). Prior research has shown that listeners integrate information across dimensions when interpreting the location of an intonational boundary (Streeter 1978, Beach 1991, de Pijper and Sanderman 1994), which suggests that perceptual weights for such boundaries may also shift in response to context. The same may also be true for lexical tones. Although pitch is the acoustic dimension which best distinguishes between the four lexical tones of Mandarin, other cues such as amplitude and duration can be used to distinguish between tones when pitch information is unavailable, as in whispered speech (Lin and Repp 1989, Blicher, Diehl & Cohen 1990, Whalen and Xu 1992, Fu and Zeng 2000, Liu and Samuel 2004; Zhang et al., under review). We would predict that creation of an artificial accent in which pitch versus durational information supported contrasting lexical tone interpretations would lead to short-term decreases in durational weighting that parallel those reported in the current paper.

Here we find that when covariation between F0 and duration is opposite that of the typical relationship in English, listeners down-weight duration but continue to rely on F0, due to F0 being a stronger cue to the presence of word emphasis in English. However, F0’s dominance as a cue to word emphasis may not be universal across all English listeners, but instead may vary as a function of the experienced overlap in the distribution of cues associated with emphasized versus not emphasized words (Holt & Lotto, 2006; Toscano and McMurray 2010). For example, individuals with congenital amusia, who have difficulty perceiving and remembering pitch in both musical and speech stimuli, weight F0 and duration roughly equally in a word emphasis categorization task like the one studied here (Jasmin et al., 2020a). We predict, therefore, that individuals with amusia would not down-weight duration when exposed to “accented” speech in which pitch and duration suggest conflicting interpretations regarding word emphasis.

Our results suggest that suprasegmental perceptual weights are not set in stone, but that they continually adjust in response to short-term speech input regularities. Future work could investigate whether suprasegmental dimensional weighting reflects the relative utility of different cues in particular listening environments, or as a function of task (Holt & Lotto, 2006). For example, prior work in segmental perception shows that weighting of an F0-based cue to voicing (the F0 of the vowel following the consonant) is increased when speech is presented in masking noise, while the duration-based cue (VOT) is down-weighted (Winn et al., 2013; Holt et al., 2018; Wu & Holt, under review). Similarly, distinct contexts and task demands are likely to impact the relative effectiveness of multidimensional acoustic information signaling suprasegmental categories, as they do for segmental categories.

Our results suggest that dimensional weighting in suprasegmental speech perception is a dynamic process, in that relative weighting can change over the time scale of just a few minutes. What neural mechanism might make possible these rapid changes in how perceptual information is integrated? One possibility is that short-term modulations in neural functional connectivity between perceptual regions devoted to the processing of a given acoustic dimension and regions associated with language processing drive changes in dimensional weighting. We recently presented evidence suggesting that functional connectivity patterns may underlie relative perceptual weighting of acoustic dimensions during suprasegmental speech perception as well (Jasmin et al. 2020d). We found that, when participants underwent fMRI scanning while performing an intonational phrase boundary perception task, connectivity between pitch-sensitive areas in the insula and superior temporal gyrus and left prefrontal language-related regions was weakened in participants with amusia, who down-weighted pitch information during suprasegmental categorization, relative to control participants. This connectivity pattern, however, could reflect intrinsic differences between amusics and controls rather than perceptual weighting. The hypothesis that dimensional weighting is linked to changes in the degree of correlated activity between task-related brain areas could be more stringently tested using the suprasegmental dimensional weighting shift paradigm presented in the current paper, by inducing shifts in cue weighting driven by contextual changes in the correlations between dimensions and examining the effects on functional connectivity. There is also important work to be done to understand which mental representations are impacted, and how distributions of speech input interact with a system tuned to expect specific regularities characteristic of a language community.

It is also possible that dimension-based statistical learning of prosodic categories may extend to production. In a study on segmental speech, exposure to a reverse (‘accented’) correlation between F0 and VOT led to down-weighting of F0 in perceptual category decisions and also diminished participants’ own use of F0 in their speech productions (Lehet & Holt, 2017). Further work could investigate whether the down-weighting of duration observed here during prosody perception also manifests in speech production acoustics, which would suggest that prosodic categories activated during perception are shared with production.

## Acknowledgments

We would like to thank Christi Gomez for her role in collecting the data. Funding: The work was funded by a Leverhulme Trust Research Project Grant (RPG-2019-107) to AT, a Leverhulme Trust Early Career Fellowship (ECF-2017-151) to KJ, and support from the National Institutes of Health (R01DC017734) to LH.

## Open Practices Statement

The data are available at BiRON https://eprints.bbk.ac.uk/id/eprint/45770 and the study was not pre-registered.

## References

Aylett, M., & Turk, A. (2004). The smooth signal redundancy hypothesis: A functional explanation for relationships between redundancy, prosodic prominence, and duration in spontaneous speech. Language and speech, 47(1), 31–56.

Bates D, Mächler M, Bolker B, Walker S (2015). “Fitting Linear Mixed-Effects Models Using lme4.” Journal of Statistical Software, 67(1), 1–48. doi: 10.18637/jss.v067.i01.

Baumann, S., Grice, M., & Steindamm, S. (2006, May). Prosodic marking of focus domains-categorical or gradient. In Proceedings of speech prosody (pp. 301–304).

Beach C (1991) The interpretation of prosodic patterns at points of syntactic structure ambiguity: evidence for cue trading relations. Journal of Memory and Language 30, 644–663.

Blicher, D. L., Diehl, R. L., & Cohen, L. B. (1990). Effects of syllable duration on the perception of the Mandarin Tone 2/Tone 3 distinction: Evidence of auditory enhancement. Journal of Phonetics, 18(1), 37–49.

Braun, B., Kochanski, G., Grabe, E., & Rosner, B. S. (2006). Evidence for attractors in English intonation. The Journal of the Acoustical Society of America, 119(6), 4006–4015.

Breen, M., Fedorenko, E., Wagner, M., & Gibson, E. (2010). Acoustic correlates of information structure. Language and cognitive processes, 25(7-9), 1044–1098.

Choi J, Hasegawa-Johnson M, Cole J (2005) Finding intonational boundaries using acoustic cues related to the voice source. JASA 118, 2579–2587.

Chrabaszcz, A., Winn, M., Lin, C. Y., & Idsardi, W. J. (2014). Acoustic cues to perception of word stress by English, Mandarin, and Russian speakers. Journal of speech, language, and hearing research, 57(4), 1468–1479.

Cole, J., Mo, Y., & Hasegawa-Johnson, M. (2010). Signal-based and expectation-based factors in the perception of prosodic prominence. Laboratory Phonology, 1(2), 425–452.

Cumming, R. E. (2010). The interdependence of tonal and durational cues in the perception of rhythmic groups. Phonetica, 67(4), 219–242.

De Pijper, J. R., & Sanderman, A. A. (1994). On the perceptual strength of prosodic boundaries and its relation to suprasegmental cues. The Journal of the Acoustical Society of America, 96(4), 2037–2047.

Dilley, L. C. (2010). Pitch range variation in English tonal contrasts: Continuous or categorical?. Phonetica, 67(1-2), 63–81.

Falé, I., & Faria, I. H. (2006, May). Categorical perception of intonational contrasts in European Portuguese. In Proceedings of Speech Prosody (pp. 69–72). Dresden, Germany: TUDpress Verlag der Wissenschaften GmbH.

Fu, Q. J., & Zeng, F. G. (2000). Identification of temporal envelope cues in Chinese tone recognition. Asia Pacific Journal of Speech, Language and Hearing, 5(1), 45–57.

Guediche, S., Blumstein, S., Fiez, J., & Holt, L. L. (2014). Speech perception under adverse conditions: insights from behavioral, computational, and neuroscience research. Frontiers in Systems Neuroscience, 7, 126.

Harmon, Z., Idemaru, K., & Kapatsinski, V. (2019). Learning mechanisms in cue reweighting. Cognition, 189, 76–88.

Holt, L. L., & Lotto, A. J. (2006). Cue weighting in auditory categorization: Implications for first and second language acquisition. The Journal of the Acoustical Society of America, 119(5), 3059–3071.

Holt, L. L., & Lotto, A. J. (2008). Speech perception within an auditory cognitive science framework. Current directions in psychological science, 17(1), 42–46.

Holt, L. L., Tierney, A. T., Guerra, G., Laffere, A., & Dick, F. (2018). Dimension-selective attention as a possible driver of dynamic, context-dependent re-weighting in speech processing. Hearing research, 366, 50–64.

Idemaru, K., & Holt, L. L. (2011). Word recognition reflects dimension-based statistical learning. Journal of Experimental Psychology: Human Perception and Performance, 37(6), 1939.

Idemaru, K., Holt, L. L., & Seltman, H. (2012). Individual differences in cue weights are stable across time: The case of Japanese stop lengths. The Journal of the Acoustical Society of America, 132(6), 3950–3964.

Idemaru, K., & Holt, L. L. (2014). Specificity of dimension-based statistical learning in word recognition. Journal of Experimental Psychology: Human Perception and Performance, 40(3), 1009.

Jasmin, K., Dick, F., Holt, L. L., & Tierney, A. (2020a). Tailored perception: Individuals’ speech and music perception strategies fit their perceptual abilities. Journal of Experimental Psychology: General, 149(5), 914.

Jasmin, K., Dick, F., & Tierney, A. T. (2020b). The Multidimensional Battery of Prosody Perception (MBOPP). Wellcome Open Research, 5(4), 4.

Jasmin, K., Sun, H., & Tierney, A. T. (2020c). Effects of language experience on domain-general perceptual strategies. Cognition.

Jasmin, K., Dick, F., Stewart, L., & Tierney, A. T. (2020d). Altered functional connectivity during speech perception in congenital amusia. Elife, 9, e53539.

Kawahara, H., & Irino, T. (2005). Underlying principles of a high-quality speech manipulation system STRAIGHT and its application to speech segregation. In Speech separation by humans and machines (pp. 167-180). Springer, Boston, MA.

Kim, D., Clayards, M., & Kong, E. J. (2020). Individual differences in perceptual adaptation to unfamiliar phonetic categories. Journal of Phonetics, 81, 100984.

Kimball, A. & Cole, J. Perception and memory for within-category detail of phonemes and pitch accents. Preprint on ResearchGate DOI: 10.13140/RG.2.2.27341.79841

Kohler, K. J. (1987). Categorical pitch perception. In Proceedings of the XIth International Congress of Phonetic Sciences (Vol. 5, pp. 331–333). Tallinn: Academy of Sciences of the Estonian Soviet Socialist Republic.

Kurumada, C., Brown, M., Bibyk, S., Pontillo, D. F., & Tanenhaus, M. K. (2014). Is it or isn’t it: Listeners make rapid use of prosody to infer speaker meanings. Cognition, 133(2), 335–342.

Kurumada, C., Brown, M., & Tanenhaus, M. K. (2018). Effects of distributional information on categorization of prosodic contours. Psychonomic bulletin & review, 25(3), 1153–1160.

Ladd, DR. (1996) Intonational phonology. Cambridge University Press.

Ladd, D. R., & Morton, R. (1997). The perception of intonational emphasis: continuous or categorical?. Journal of phonetics, 25(3), 313–342.

Lehet, M., & Holt, L. L. (2017). Dimension-based statistical learning affects both speech perception and production. Cognitive science, 41, 885–912.

Lisker, L. (1957). Closure duration and the intervocalic voiced-voiceless distinction in English. Language, 33(1), 42–49.

Lisker, L. (1986). “Voicing” in English: A catalogue of acoustic features signaling/b/versus/p/in trochees. Language and speech, 29(1), 3–11.

Liu, R., & Holt, L. L. (2015). Dimension-based statistical learning of vowels. Journal of Experimental Psychology: Human Perception and Performance, 41(6), 1783.

Liu, S., & Samuel, A. G. (2004). Perception of Mandarin lexical tones when F0 information is neutralized. Language and speech, 47(2), 109–138.

Nath, A. R., & Beauchamp, M. S. (2011). Dynamic changes in superior temporal sulcus connectivity during perception of noisy audiovisual speech. Journal of Neuroscience, 31(5), 1704–1714.

Peppé, S., Maxim, J., & Wells, B. (2000). Prosodic variation in southern British English. Language and Speech, 43(3), 309–334.

Pierrehumbert, J., & Hirschberg, J. B. (1990). The meaning of intonational contours in the interpretation of discourse.

Pierrehumbert, J. B., & Steele, S. A. (1989). Categories of tonal alignment in English. Phonetica, 46(4), 181–196.

Remijsen, B., & van Heuven, V. J. (1999). Gradient and categorical pitch dimensions in Dutch: diagnostic test. In Proceedings of the 14th International Congress of Phonetic Sciences (Vol. 2, pp. 1865–1868).

Repp, B. H., & Lin, H. B. (1989). Acoustic properties and perception of stop consonant release transients. The Journal of the Acoustical Society of America, 85(1), 379–396.

Roettger T, Franke M (2019) Evidential strength of intonational cues and rational adaptation to (un-) reliable intonation. Cognitive Science 43, e12745.

Roettger, T. B., & Rimland, K. (2020). Listeners’ adaptation to unreliable intonation is speaker-sensitive. Cognition, 204, 104372.

Rohe, T., & Noppeney, U. (2018). Reliability-weighted integration of audiovisual signals can be modulated by top-down attention. eneuro, 5(1).

Russell V. Lenth (2016). Least-Squares Means: The R Package lsmeans. Journal of Statistical Software, 69(1), 1-33. <doi:10.18637/jss.v069.i01>

Saindon, M. R., Trehub, S. E., Schellenberg, E. G., & van Lieshout, P. H. (2017). When is a Question a Question for Children and Adults?. Language Learning and Development, 13(3), 274–285.

Schertz, J., Cho, T., Lotto, A., & Warner, N. (2016). Individual differences in perceptual adaptability of foreign sound categories. Attention, Perception, & Psychophysics, 78(1), 355–367.

Saindon M, Cirelli L, Schellenberg E, van Lieshout P, Trehub S (2017) Children’s and adults’ perception of questions and statements from terminal fundamental frequency contours. JASA 141, 3123–3131.

Schneider, K., & Lintfert, B. (2003, August). Categorical perception of boundary tones in German. In Proceedings of the 15th International Conference of the Phonetic Sciences (pp. 631–634).

Streeter, L. A. (1978). Acoustic determinants of phrase boundary perception. The Journal of the Acoustical Society of America, 64(6), 1582–1592.

Toscano, J. C., & McMurray, B. (2010). Cue integration with categories: Weighting acoustic cues in speech using unsupervised learning and distributional statistics. Cognitive science, 34(3), 434–464.

Turk, A. E., & Sawusch, J. R. (1997). The domain of accentual lengthening in American English. Journal of Phonetics, 25(1), 25–41.

Turk, A. E., & White, L. (1999). Structural influences on accentual lengthening in English. Journal of phonetics, 27(2), 171–206.

Watson, D. G., Tanenhaus, M. K., & Gunlogson, C. A. (2008a). Interpreting pitch accents in online comprehension: H* vs. L+ H. Cognitive science, 32(7), 1232–1244.

Watson, D. G., Arnold, J. E., & Tanenhaus, M. K. (2008b). Tic Tac TOE: Effects of predictability and importance on acoustic prominence in language production. Cognition, 106(3), 1548–1557.

Watson, D. G. (2010). The many roads to prominence: Understanding emphasis in conversation. In Psychology of learning and motivation (Vol. 52, pp. 163–183). Academic Press.

Whalen, D. H., & Xu, Y. (1992). Information for Mandarin tones in the amplitude contour and in brief segments. Phonetica, 49(1), 25–47.

Winn, M. B., Chatterjee, M., & Idsardi, W. J. (2013). Roles of voice onset time and F0 in stop consonant voicing perception: Effects of masking noise and low-pass filtering. Journal of Speech, Language, and Hearing Research.

Wu, Y., & Holt, L. L. (2018). Phonetic category activation drives dimension-based adaptive tuning in speech perception. In CogSci.

Wu, Y.C. (2020). Behavioral, computation, and electrophysiological investigations of adaptive plasticity mechanisms in speech perception. Doctoral Dissertation, Carnegie Mellon University. ProQuest Dissertations & Theses Global.

Wu, Y. C., & Holt, L. L. (under review). Category activation drives adaptive plasticity in dimension-based statistical learning in speech perception.

Xu, Y., & Xu, C. X. (2005). Phonetic realization of focus in English declarative intonation. Journal of Phonetics, 33(2), 159–197.

Yang, X., Shen, X., Li, W., & Yang, Y. (2014). How listeners weight acoustic cues to intonational phrase boundaries. PloS one, 9(7), e102166.

Zárate-Sández, G. (2016). Categorical perception and prenuclear pitch peak alignment in Spanish. Proceedings of Speech Prosody 2016, 663–667.

Zhang, X., & Holt, L. L. (2018). Simultaneous tracking of coevolving distributional regularities in speech. Journal of Experimental Psychology: Human Perception and Performance, 44(11), 1760.

Zhang, X., Wu, X., & Holt, L. L. (2021). The learning signal in perceptual tuning of speech: Bottom-up vs. Top-down information. Cognitive Science 45, 312947

Zhang, H., Wiener, S., & Holt, L. L. (under review). Evidence for dynamic adjustment of cue weighting in speech.

